# Specialization of a polyphenism switch gene following serial duplications in *Pristionchus* nematodes

**DOI:** 10.1101/055384

**Authors:** Erik J. Ragsdale, Nicholas A. Ivers

## Abstract

Polyphenism is an extreme manifestation of developmental plasticity, requiring distinct developmental programs and the addition of a switch mechanism. Because the genetic basis of polyphenism switches has only begun to be understood, how their mechanisms arise is unclear. In the nematode *Pristionchus pacificus*, which has a mouthpart polyphenism specialized for alternative diets, a gene (*eud-1*) executing the polyphenism switch was recently identified as the product of lineage-specific duplications. Here we infer the role of gene duplications in producing a switch gene. Using reverse genetics and population genetic analyses, we examine evidence for competing scenarios of degeneration and complementation, neutral evolution, and functional specialization. Of the daughter genes, *eud-1* alone has assumed switch-like regulation of the mouth polyphenism. Measurements of life-history traits in single, double, and triple sulfatase mutants did not, given modest sample sizes and a benign environment, identify alternative or complementary roles for *eud-1* paralogs. Although possible roles are still unknown, selection analyses of the sister species and 104 natural isolates of *P. pacificus* detected purifying selection on the genes, suggesting their functionality by their fixation and evolutionary maintenance. Our approach shows the tractability of reverse genetics in a non-traditional model system to study evolution by gene duplication.

---

Developmental polyphenism, or the production of discrete alternative phenotypes from a single genotype, can afford an immediate adaptive response to disparate environmental problems. Developmental plasticity in general is considerably widespread among multicellular organisms (Pigliucci 2001; West-Eberhard 2003; Pfennig et al. 2010), although polyphenism is much less common, often simply the result of discontinuities in the environment (Nijhout 2003). In contrast to norms of reaction, true polyphenism requires distinct developmental programs, each with their own fitness advantages (Moran 1992), and a switch mechanism. The question of how polyphenism ultimately arises has garnered some attention, and alternative morphs have been inferred or shown to arise from conditionally expressed variation, especially in a context of pre-existing plasticity (Nijhout 2003; Suzuki and Nijhout 2006; Ledón-Rettig et al. 2010; Rajakumar et al.2013; Susoy et al. 2016). However, the genetic origins of polyphenism are still unclear.In particular, how switch mechanisms for polyphenisms appear and become accommodated is unknown, and how genes execute them was until recently hypothetical.

In the dimorphic nematode *Pristionchus pacificus*, a morphological polyphenism was found to operate through a switch gene (Ragsdale et al. 2013). In this species and others in the family Diplogastridae, one of two alternative adult morphs develops irreversibly in response to environmental cues. In the presence of resource stresses including crowding and starvation, a “eurystomatous” (Eu) morph with predatory mouthparts develops (Bento et al. 2010; Bose et al. 2012; Serobyan et al. 2013), allowing a shift from the ancestral diet of microbes, for which the alternative “stenostomatous” (St) morph is specialized (Serobyan et al. 2014; Wilecki et al. 2015). In *P. pacificus* the two morphs differ qualitatively in form, including the presence of an additional tooth and other mouth armature (Ragsdale 2015). The identified switch gene encodes the sulfatase EUD-1, which was hypothesized to modify an unknown hormone based on the presence of sulfated sterols in *Caenorhabditis elegans* (Carroll et al. 2006; Hattori et al. 2006). The gene is one of three duplicate homologs of a single sulfatase-encoding gene, *sul-2,* in outgroups that lack the mouth polyphenism, including *C. elegans* (Fig. 1). However, the ancestral role of SUL-2 is unknown, as no function for the gene in *C. elegans* has been described. Because *eud-1* was the only haploinsufficient gene identified in a forward screen for *P. pacificus* polyphenism mutants (Ragsdale et al. 2013), it was unclear whether similarly acting genes, such as close paralogs, could inform the developmental logic the switch. Furthermore, the role of gene duplications in the evolution of *eud-1* as a polyphenism regulator was previously unknown.

**Figure 1.**
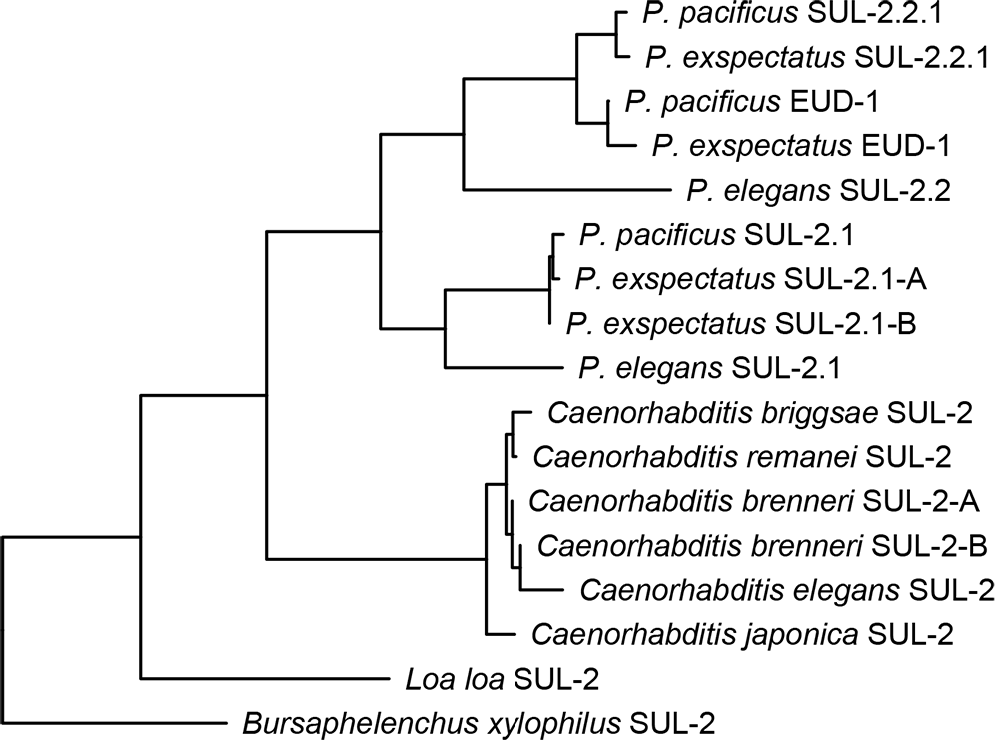
Phylogenetic history of *sul-2* homologs in *Pristionchus* and other nematodes. *sul-2,* which encodes an arylsulfatase, has undergone serial duplications in the lineage of *P. pacificus* to produce the polyphenism switch gene *eud-1.* Recent duplications of *sul-2* homologs have also occurred in other lineages, including those of *P. exspectatus* and *Caenorhabditis brenneri.* Tree was inferred from protein sequences and is summarized from Ragsdale et al. (2013).

Gene duplication has long been thought to allow the evolution of functional novelty (Haldane 1932; Muller 1940; Ohno 1970), and examples of such novelty have been thoroughly attested by empirical studies (e.g., True and Carroll 2002; Hurley et al. 2005; Irish and Litt 2005). Because the gene regulating the *P. pacificus* polyphenism is also one of several serially duplicated genes, analysis of paralogous genes might reveal how genomic processes such as gene duplication affect the function or specialization of a plasticity switch. For example, the persistence of an ancestral function as a polyphenism regulator among paralogs would support a scenario of selection for gene amplification in this species (Bergthorsson et al. 2007). Alternatively, duplicated genes may have become partially or completely specialized, whether for complementary functions (Force et al. 1999) or by release of a pleiotropic gene from adaptive conflict (Hughes 1994; Hittinger and Carroll 2007). Another possibility is that duplications of a pre-existing switch gene may simply be neutral genomic processes, as would be detectable by apparent nonfunctionalization of recent duplicates (Lynch and Conery 2000). A combination of functional tests with inferences of molecular evolution could distinguish among these alternative scenarios.

Two features of *P. pacificus* as an emerging model system promote the functional study of evolution by gene duplication. First, extensive genomic resources are available for *P. pacificus* (Dieterich et al. 2008) that allow detection of all recognizable paralogs of a given gene. The second is tested techniques for germline transgenesis (Schlager et al. 2009). Building upon these resources, the use of CRISPR/Cas9 to induce target-specific gene knockouts in *P. pacificus* (Lo et al. 2013; Witte et al. 2015) has suggested the ability to link phenotypes to genes of unknown function. Furthermore, the availability of genomic sequence variants for its close sibling species *P. exspectatus* (Kanzaki et al. 2012) and over 100 isolates of *P. pacificus* allows tests for signatures of selection on duplicated genes across populations (Rödelsperger et al. 2014). Here, we have used reverse genetics and analyses of natural variation to investigate the function of all *sul-2* duplicates in *P. pacificus.* As a result, we show that polyphenism regulation has evolved by the specialization of a single control gene following serial duplications.

## Materials and Methods

### Reverse genetics

To determine the phenotype of previously uncharacterized homologs of *Cel-sul-2* in *P. pacificus,* namely *Ppa-sul-2.1* and *Ppa-sul-2.2.1,* we generated mutant lines for phenotypic assays. We induced mutations in these genes using the type-II CRISPR/Cas9 system (Jinek et al. 2012), consistent with a protocol recently published for *P. pacificus* (Witte et al. 2015). In summary, we targeted the germline of wild-type (PS312, “California”) *P. pacificus* with a cocktail including Cas9 caspase protein and a universal *trans*- activating CRISPR RNA ligated to custom 23‐bp single guide RNAs (sgRNAs), as synthesized by ToolGen, Inc. (Seoul, Korea). The transformation cocktail was first mixed from: 2 µl Cas9 protein at 5 µg/µl in water; 2µl protein storage buffer (20 mM HEPES-KOH pH 7.5, 150 mM KCl, 1 mMDTT); 0.5 µl each of two sgRNAs with unique target regions (see below) and each stored at 6 µg/µl in protein storage buffer. This mix was incubated for 37 °C for 10 minutes, followed by a 10‐fold dilution to 50 µl in TE buffer (10 mM Tris-HCI, 1 mM EDTA, pH 8.0). After centrifuging the cocktail at 4 °C for 10 minutes to separate any precipitates, the solution was loaded into a needle pulled from a 1.0-mm diameter borosilicate glass capillary. We delivered the cocktail into both ovaries of adult hermaphrodites (i.e., self-fertilizing, morphological females) by microinjection, using the method described for DNA-mediated transformation of *P. pacificus* (Schlager et al. 2009). We performed injections on a Zeiss Axiovert coupled to an Eppendorf TransferMan micromanipualtor and Eppendorf FemtoJet injector.

For each transformation, we introduced two different sgRNAs to ensure the success of at least one. sgRNAs were designed to target an exon that encodes a conserved active site (Waldow et al. 1999). Whereas sgRNAs for *sul-2.1* were gene-specific, sgRNAs designed against *sul-2.2.1* and *eud-1* were identical, making it possible to create single and double mutants with the same CRISPR injection. An sgRNA specific to two genes was necessary to generate double mutants, which due to the close physical linkage of the two genes (ca. 6 kb) would be infeasible to create by crossing separate mutant lines. sgRNA sequences did not match any part of the *P. pacificus* genome other than their target sequences.

Immediately after injection, P0 hermaphrodites were isolated onto individual culture plates for self-crossing. Because of the anticipated low success rate of injections (Schlager et al. 2009), the chances of two independently mutated gametes fusing were presumed to be small, such that any mutant F1 were likely to be complemented by a wild-type allele at the targeted locus. We therefore established lines homozygous for mutant alleles by cloning the F2 of injected hermaphrodites by allowing F1 isolated as virgins, namely at the pre-adult (J4) stage, to self-cross. After leaving F2, we sacrificed F1 adults to determine their genotype at the region of the predicted lesions. To detect the lesions, we designed a pair of amplification primers that were in combination unique for each gene (Table S1). Because of the high similarity of sequences in and near the targeted gene region of *sul-2.2.1* and *eud-1,* the reverse primers were identical, but in combination with distinct forward primers the genes were selectively amplified. Sequencing reactions were performed using forward primers. If an F1 genotype was heterozygous for a frame-shift mutation, which was detected by the presence of two conflicting sequences 3’ of the predicted lesion site in the sequence chromatogram, 30 F2 from that individual were isolated. After letting the F2 produce F3, we sacrificed the F2 to identify those that were homozygous for each mutation and thus a clonal mutant line. We then confirmed homozygosity of the mutation by genotyping 30 F3 from the established mutant line. Using this method, we could detect genetic lesions and confirm the homozygosity of mutant lines blind to possible mutant phenotypes. For lines transformed with the nonspecific sgRNA, we genotyped both *sul-2.2.1* and *eud-1.*

Upon generating mutant lines, we created lines carrying multiple mutations to detect complementary function (i.e., subfunctionalization) of genes for assayed phenotypes. We used crosses, together with the simultaneous creation of a double mutant by CRISPR/Cas9, to create a *sul-2.1; sul-2.2.1* double mutant and a *sul-2.1; sul-2.2.1 eud-1* triple mutant. We confirmed homozygosity of mutant genes in double and triple mutants blind to phenotype by sequencing all three loci of 10 individuals per clonal line. Chromatograms for mutant sequences have been archived in the Dryad Digital Repository (datadryad.org).

### Phenotypic assays

We assayed the polyphenism and standard life-history traits in all single, double, and triple mutants above as well as the wild-type strain PS312 and a previously generated line carrying a null allele *(tu445)* for *eud-1.* Because of the high homozygosity of the inbred wild-type strain, all lines except for that carrying *eud-1(tu445),* which was previously produced by chemical mutagenesis and backcrossed twice (Ragsdale et al. 2013), were presumed to have identical genetic backgrounds to the mutated genes.

Prior to assays, we took several measures to prevent any unknown epigenetic effects on phenotypes (Rechavi et al. 2011). First, all lines were well fed on monoxenic food source *(Escherichia coli* OP50) in a standard amount (lawns grown from 300 µl L-broth) for at least five generations at ~23 °C. To control for population densities as far as possible without knowing differences in mutant growth rates a priori, each generation was established from five virgin hermaphrodites. To minimize potential variation of pheromone concentrations, all cultures were kept on a 6-cm diameter plate supplied with 10 ml of nematode growth medium (NGM) agar.

We assayed the effect of mutant genes on the mouth polyphenism by recording the ratio of morphs in the wild type and all combinations of mutants. Specifically, we scored this ratio as the frequency of Eu individuals born of a single virgin hermaphrodite under the culture conditions described above. All individuals that were not Eu were St, except for rare cases (<0.5%) of apparent intermediates, which were excluded from counts. We determined mouth phenotypes by differential interference contrast (DIC) microscopy on a Zeiss Axioscope, referring to a suite of qualitative morphological differences described previously (Serobyan et al. 2013). Up to 80 individuals on each of 9-11 replicate plates (N = 558−711) were screened for each line, except for those lines with a mutant allele of *eud-1,* where 3 plates (N = 240) were screened per line. We started all polyphenism experiments from a single cohort (stock plate).

To explore whether any *sul-2* paralogs have other, major phenotypic consequences, we measured the effects of deletion mutants on two basic life-history traits. Specifically, we recorded brood sizes and growth rates for the wild-type strain and all single, double, and triple mutants. The first of these traits, lifetime fecundity, was chosen based on its use as a standard life-history measure (Gilarte et al. 2015) and its demonstrated ability to detect fitness differences on alternative diets in *P. pacificus* (Rae et al. 2008; Serobyan et al. 2014). We measured the second trait, growth rate, because of the previous detection of differences in rates between the two mouth-morphs on a similar diet, albeit with a finer time-course than what we employ here (Serobyan et al. 2013). For each line, we isolated multiple mid-to late-J4 individuals of a single cohort from our well-fed, density-controlled stocks. Single hermaphrodites then developed to maturity while feeding ad libitum on OP50 as described above. Starting at 48 hours after transfer onto individual plates, all adult offspring were counted and removed every 12 hours for 3.5 days, at which point all offspring, with the exception of a single replicate of one treatment, had reached adulthood. In this manner we monitored the cumulative proportion of broods completing development over time. Together, these measurements recorded the fecundity of the parents and rates of development. In addition to this experiment, which we performed in six replicates (i.e., from six transferred J4) per line, we continued to measure brood sizes for additional replicates from these cohorts and from cohorts of three further generations. Specifically, we counted hatched, viable offspring three days after isolating their mother as a J4. In contrast to simply counting eggs, our assay captured absolute fecundity, which inherently includes other effects on fitness: fertilization by self-sperm, the production of which is also a sensitive measure of fitness (Serobyan et al. 2014), and survivorship of larvae through early postembryonic development when feeding ad libitum at low population densities. In total, we recorded brood sizes for a total of *n* = 39−45 mothers for each line. All raw, quantitative phenotypic data for fitness measures and the mouth polyphenism have been archived in the Dryad repository.

In addition to life-history traits of directly developing individuals, we determined whether any of the mutants, either singly or in combination, were dauer-formation-defective (Riddle et al. 1981). The dauer stage is a metabolically quiescent, dispersal stage that is a shunt to an intermediate, direct-developing larval (J3) stage. Like the Eu morph, the dauer develops in response to crowding or low-food conditions and shares some developmental regulation with the mouth polyphenism in *P. pacificus* (Ogawa et al. 2009; Bento et al. 2010). The dauer is morphologically distinct from directly developing larval stages by having a slim body, a closed mouth, enlarged amphid openings, and extensive wax droplets at the body surface. We assayed whether lines could produce dauers in respond to starvation, specifically by letting populations exhaust their food source. Dauer induction was scored as presence or absence. To induce dauers, we started cultures from five J4 hermaphrodites on plates prepared as above and keeping them on those plates without further feeding for four weeks. Experimental plates were screened for dauers using a Zeiss Discovery V20 stereomicroscope, supplemented by DIC microscopy.

### Selection analyses

As an alternative test of function of *eud-1* paralogs in *P. pacificus,* we looked for evidence of selection on these genes. Specifically, we tested the alternative possibilities of purifying (negative) selection or neutral evolution of *eud-1* paralogs. The former case would suggest evolutionary maintenance of a given gene, whether or not its function could be identified, and the latter that the duplicates were in the process of becoming pseudogenes. For analysis, we used the complete predicted coding regions of *sul-2.1* and *sul-2.2.1,* previously inferred with AUGUSTUS (Stanke et al. 2004; Rödelsperger et al. 2014; www.pristionchus.org). We extracted the following archived genomic sequences: Contig65-snap.11, Contig8-snap.31, which represent *Ppa-sul-2.1* and *Ppa-sul-2.2.1,* respectively; scaffold22-pex_aug2013-10597.t1, representing *Pex-sul-2.2.1; eud-1* of both species, as informed by RACE of *Ppa-eud-1* (Ragsdale et al. 2013). Extracted sequences belonged to 104 haplotypes of *P. pacificus* and one haplotype of *P. exspectatus,* strain RS5522B (Rödelsperger et al. 2014). Because *sul-2.1* has two homologs in *P. exspectatus* (Ragsdale et al. 2013), *P. exspectatus* was not included in selection analyses of *sul-2.1.* All genes were previously confirmed in part by expressed mRNAs. For parts of the genes that have not yet been confirmed, we used predicted exons. Based on high sequence similarity of open reading frames and splice sites between *Pex-sul-2.2.1* with *Ppa-sul-2.2.1,* we identified two putative exons of the latter in the raw genome sequence additional to the AUGUSTUS annotation. Genes were aligned automatically by codon in MUSCLE (Edgar 2004) and then manually in MEGA 7 (Tamura et al. 2011). The result of manual alignment was to exclude the three terminal codons of *Ppa-sul-2.2.1,* which are absent in *P. exspectatus,* and the terminal codon of *Pex-eud-1,* which is absent in *P. pacificus.*

To test for patterns of selection, we estimated the ratio of rates of non-synonymous to synonymous substitutions (*p*_N_/*p*_S_) for the predicted coding regions of each gene. Substitutions were inferred using the single likelihood ancestor counting (SLAC) method, a conservative approach to detect selection in large numbers (>40) of sequences (Kosakovsky Pond and Frost 2005). The counting procedure was implemented in HyPhy on the Datamonkey web server (Delport et al. 2010). Substitution histories were each reconstructed on a gene tree inferred by maximum likelihood as implemented in RAxML 8 (Stamatakis 2006). We used the most likely tree from 100 independent analyses of a single data partition and invoking a general time reversible (GTR) model of substitution and the CAT approximation of rate heterogeneity. In the SLAC procedure, the codon model was fitted allowing a GTR model of nucleotide evolution. For each gene we estimated the mean *p*_N_/*p*_S_, for which values of <1 were interpreted to indicate purifying selection; significance was determined if a 95% profile likelihood confidence interval did not include the null expectation (*p*_N_/*p*_S_ = 1). We also tested hypotheses of selection on individual codons of each gene, with significance assessed using a two-tailed test on an extended binomial distribution. To detect macroevolutionary signatures of selection, we estimated *d*_N_/*d*_S_ for *sul-2.2.1* with respect to its homolog in *P. exspectatus.* We estimated this ratio with respect to the numerous sequenced haplotypes of *P. pacificus* by calculating the mean pairwise *d*_N_/*d*_S_ between *P. exspectatus* and all variants of *P. pacificus.* Pairwise *d*_N_/*d*_S_ was estimated using the method of Yang and Nielsen (2000) as implemented in PAML 7 (Zhang 2007). Because this inference method differed from that of *p*_N_/*p*_S_, including by the absence of phylogenetic structure, we do not compare our *d*_N_/*d*_S_ and *p*_N_/*p*_S_ estimates.

### Statistical analyses of phenotypic data

All statistical analyses of phenotypic data were conducted in R 3.2.3 (R Core Team 2013). For analysis of plasticity phenotypes, recorded as proportional data, we performed logistic regression using a binomial error structure and a logit link function. With this analysis, we specifically tested for differences among lines without a *eud-1* mutation. Replicates within lines were included in the model, and significance was assessed using a z-test. In an additional test, we pooled samples from replicates within lines and performed pairwise comparisons of mutant lines without a *eud-1* mutation to the wild type using Fisher's exact test. Significance thresholds in the latter test were set using a Bonferroni correction of α.

We evaluated differences in fecundity among lines following a Shapiro-Wilk test to determine normality. Because data for the *iub3* line, the *iub2; iub3* line, and the triple mutant line were not normally distributed (*P* > 0.05 for all), we used a Kruskal-Wallis rank sum test to compare distributions of all lines. We investigated potential individual differences to the wild-type strain using post-hoc pairwise Kruskal-Wallis tests. Lastly, to examine differences in growth rates, which were recorded as cumulative distributions, we performed two-sample Kolmogorov-Smirnov tests on each non-wild-type variable with respect to the wild type. Significance was determined for planned pairwise comparisons of fecundity and growth rates using the Bonferroni method.

## Results

### Mutants for switch-gene paralogs produced by reverse genetics

Targeted DNA editing successfully produced heritable mutations in all identifiable homologs of the sulfatase-encoding gene *sul-2* in *P. pacificus* (Fig. 2). We established lines with the following mutant alleles: *sul-2.1(iub3), sul-2.2.1(iub1), sul-2.2.1(iub2),* and *eud-1(iub14).* All mutant lines were viable when homozygous for the mutant alleles as single, double, and triple mutants. The efficiency of our transformations were on the order of 1% of examined F1: injections against *sul-2.2.1* and *eud-1* generated 2 independent mutant lines from 439 F1 screened (from 22 P0), and injections against *sul-2.1* produced 1 mutant line from 192 screened (36 P0). Genetic lesions included both deletions and random insertions, causing frame shifts in all alleles. Among the mutant lines was a simultaneous double mutant, *sul-2.2.1(iub1) eud-1(iub14).* In this line both genes showed identical mutations, possibly due to homology-directed repair (e.g., Dickinson et al. 2013) between the identical, non-orthologous sequence regions targeted. This result is reminiscent of previous observations on non-specific gene ablations, which sometimes produced identical deletions in multiple genes (Peng et al. 2015). In summary, our results show the feasibility of using CRISPR/Cas9 to produce multiple, simultaneous mutants of unknown phenotype in this nematode system.

**Figure 2.**
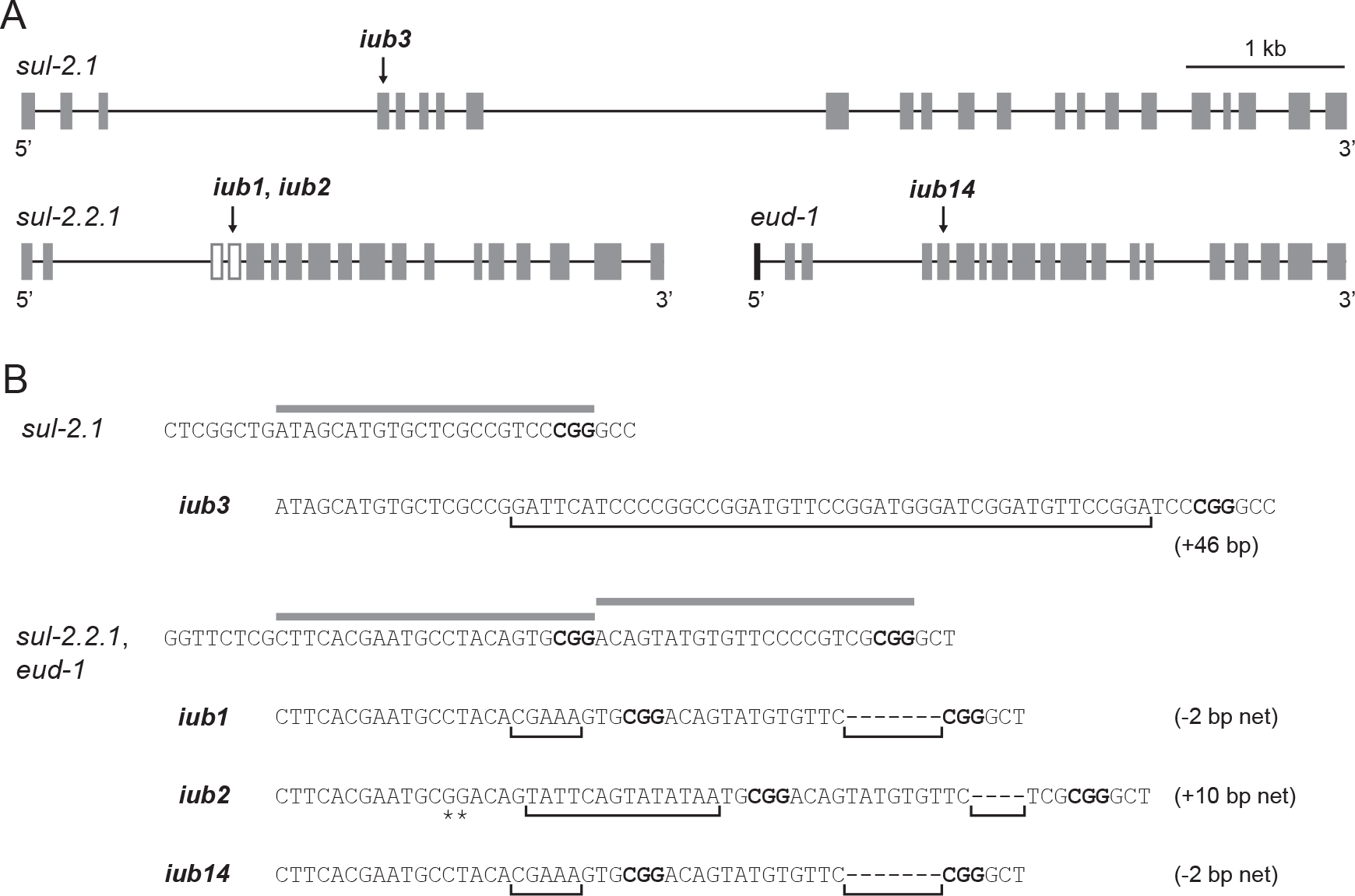
Mutant alleles of *sul-2* homologs in *P. pacificus.* Mutants were produced using type II CRISPR/Cas9. (A) Predicted or confirmed gene structure of all *sul-2* homologs, showing sites of induced genetic lesions in alleles *iubl, iub2, iub3,* and *iub14.* In all three genes, which encode arylsulfatases, target sequences encode a conserved active site. Gene structures are drawn to same scale, gray boxes representing exons. For *sul-2.2.1,* exons represented by open boxes were manually identified. Exon (black) at 5’ end of *eud-1* has no identifiable homolog upstream of *sul-2.2.1* and encodes part of a putative signal peptide, as predicted by the SignalP 4.1 server (Petersen et al. 2013). (B) Genetic lesions in mutant alleles. Gray bars, which are above wild-type sequences, indicate sgRNA sequences. *sul-2.2.1(iub1)* and *eud-1(iub14),* which were produced in the same germline, incurred identical lesions. In addition to the insertion and deletions in *sul-2.2.1(iub2),* two nearby, additional mutations (asterisks) were confirmed. Protospacer-associated motif (CGG) is in boldface font. Brackets indicate deletions and random insertions.

### A single gene duplicate controls the polyphenism switch

The ability to isolate lines homozygous for mutations in one, two, or all duplicate sulfatase genes allowed us to distinguish the effects of single genes as well as complementary roles of possibly subfunctionalized genes. We hypothesized that if the other sulfatases had any residual function in the polyphenism switch, the phenotypes of individual mutants should be apparent and their effects as multiple mutants should be additive. Measurements of the mouth-polyphenism ratio under standard environmental conditions revealed that only one gene, *eud-1,* had an effect as a single mutant (Fig. 3). This effect, as previously reported, was completely penetrant, as was its effect in combination with other mutations (Fig. 3). Furthermore, neither single-nor doublemutant lines with a wild-type copy of *eud-1* had any significant effect on the mouth polyphenism with respect to the wild type (z-test, *P* > 0.05; Fisher's exact test, α' = 0.012; Fig. 3; Table S2). This result shows that even when homozygous for null alleles, neither of the two paralogous sulfatases in *P. pacificus* have retained, if they ever had, a function analogous to that of *eud-1* in the polyphenism. Thus, both serial duplications of the ancestral sulfatase-encoding *sul-2* have been followed by the specialization of a single duplicate as a polyphenism switch gene.

**Figure 3.**
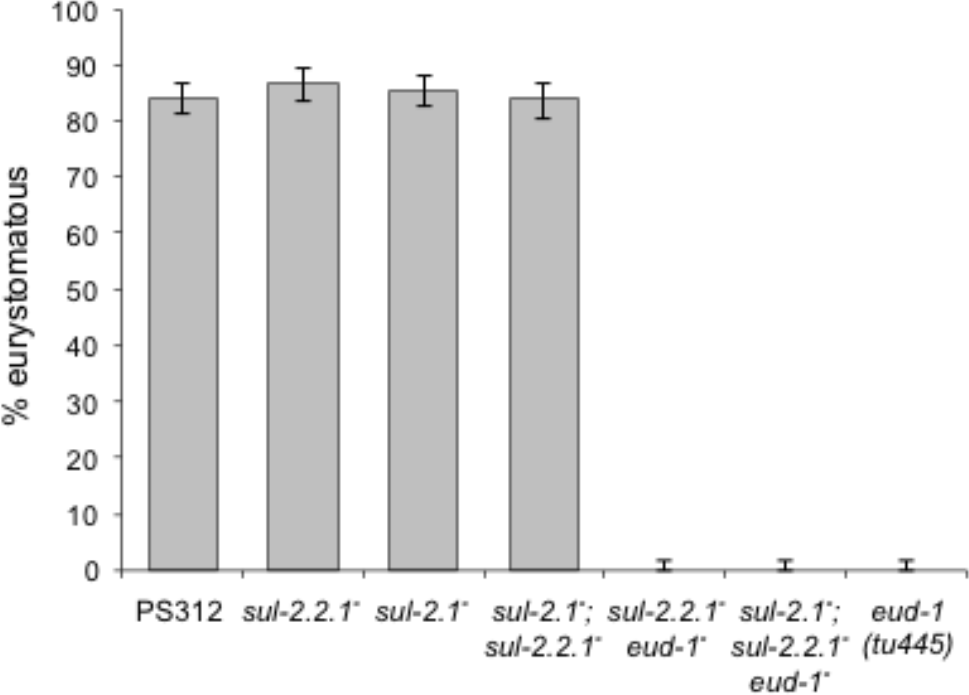
Mouth-polyphenism phenotype in wild-type and mutant hermaphrodites for *sul-2* homologs of *P. pacificus.* Frequency of the eurystomatous (Eu) form was measured under a common environmental regimen for all strains. The wild type strain is the reference strain PS312. Mutant alleles (̄) are: *sul-2(iub3), sul-2.2.1(iub1), sul-2.2.1(iub2),* and *sul-2.2.1(iub14).* Allele of *sul-2.2.1* occurring as single mutant or with only *sul-2.1* is *iub2*; allele of *sul-2.2.1* occurring with *eud-1* is *iub1.* For graphical representation, mouth-morph frequencies were pooled across replicates within lines, with confidence intervals estimated using a binomial test. These results are qualitatively identical to those derived from both the generalized linear model and Fisher's exact test with corrections for multiple comparisons as described in text.

### Alternative functions of switch-gene paralogs are unknown

In the absence of obvious qualitative phenotypes for mutant paralogs of *eud-1,* we measured two basic life-history traits to explore whether the paralogs have subfunctionalized for some other major effect in a benign environment. We hypothesized that if other functions of these genes were essential for normal development and physiology, multiple deletions of the genes would reveal differences in growth or fecundity with respect to the wild type. When we measured growth rate, no significant differences between the wild type and any single, double, or triple mutant line was detected (Kolmogorov-Smirnov, α’ = 0.009; Fig. 4A). However, fecundity differed among lines (Kruskal-Wallis, *P* < 10^-5^, *χ*^2^= 35.93, df = 6; Fig. 4B). Post-hoc pairwise comparisons with the wild type suggested that one line with a higher median, *sul-2.2.1(iub2),* had different fecundity than the wild type (P < 0.009; Fig 4B). Neither line with a lower median fecundity than the wild type differed from that strain (Table S3). The only line with a possibly lower fecundity was the double mutant *sul-2.2.1(iub1) eud-1(iub14)* (n.s., α' = 0.009), although this contrasted with the fecundity of the triple mutant, which did not differ from the wild type despite carrying the same mutations of the double mutant (Table S3). Taken together, our first screen for fitness consequences did not reveal unambiguous phenotypic defects of mutant alleles, suggesting that more nuanced phenotypic assays will be necessary to identify the functions of *sul-2.1* and *sul-2.2.1.*

**Figure 4.**
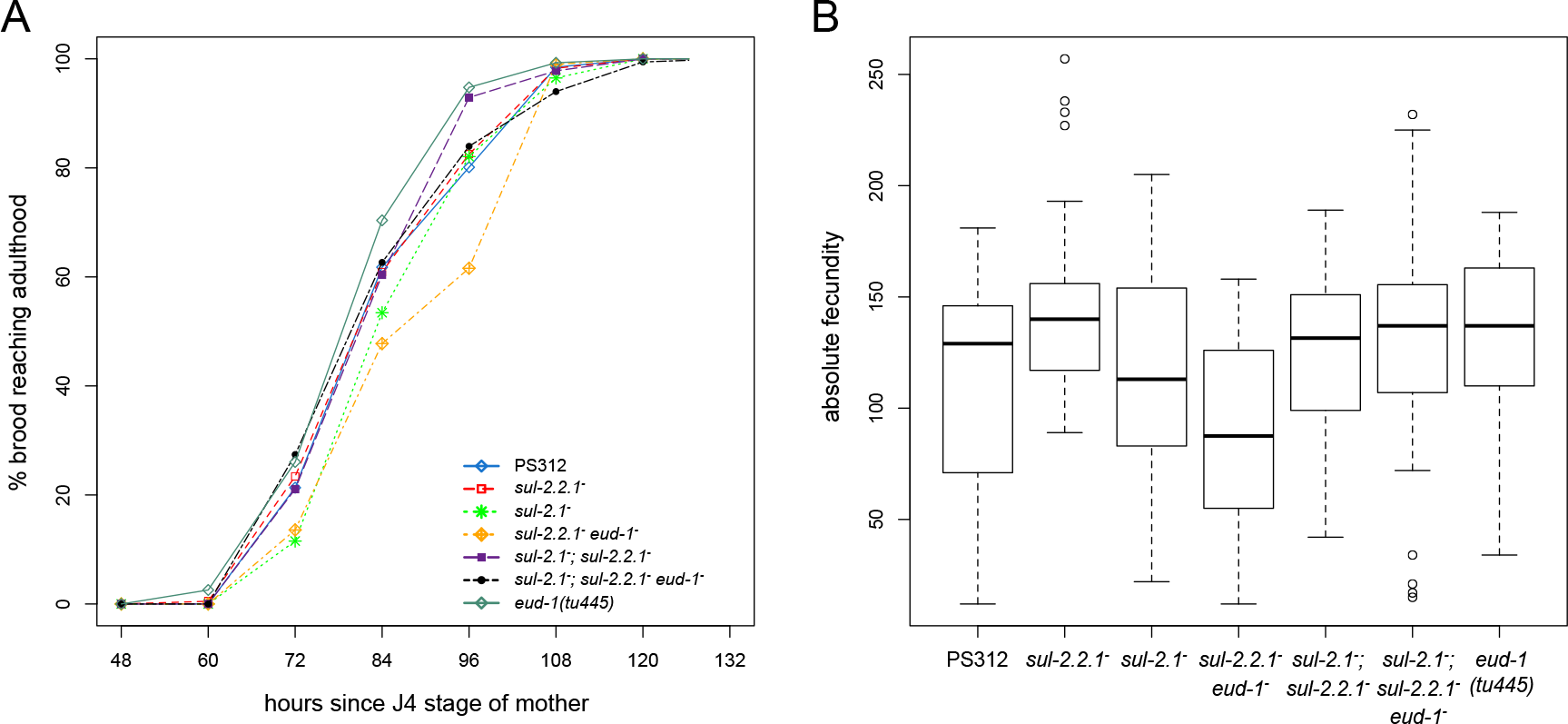
Life-history traits of the wild type and mutants for *sul-2* homologs of *P. pacificus* fed ad libitum under standard laboratory conditions. The wild type strain is the reference strain PS312. Mutant alleles (̄) are: *sul-2(iub3), sul-2.2.1(iub1), sul-2.2.1(iub2),* and *sul-2.2.1(iub14)*. Allele of *sul-2.2.1* occurring as single mutant or with only *sul-2.1* is *iub2*; allele of *sul-2.2.1* occurring with *eud-1* is *iub1.* Except for the single *eud-1* mutant *(tu445),* which was previously produced by chemical mutagenesis, the genetic backgrounds of the wild-type and mutants *sul-2.1(iub3), sul-2.2.1(iub1), sul-2.2.1(iub2),* and *eud-1(iub14)* are inbred and presumed identical. (A) Time to maturity, recorded as the cumulative percentage of offspring reaching maturity in the time since the J4 stage of the mother. Variation in time to maturity represents individual developmental rates as well as timing of oviposition by the mother. (B) Absolute fecundity, measured as the number of hatched, viable offspring per self-fertilized hermaphrodite. Box plots show the median (center bar), lower and upper quartiles (box bounds), non-outlier range limits (whiskers), and outliers (circles).

All single, double, and triple mutants were able to produce dauers (data not shown). Our brief assay showed that, despite being homozygous for null alleles, all lines readily produced dauers after exhausting bacterial food on plates. Therefore, no sulfatase mutant was fully penetrant for a dauer-formation-defective phenotype, and any involvement of the other sulfatases must be quantitative or context-dependent if present.

In summary, our results do not reveal the roles of the examined sulfatases other than EUD-1 in the mouth polyphenism switch. Furthermore, we found no evidence of degeneration and complementation for an obvious function between any two, or among all three, sulfatases in our screen for coarse fitness phenotypes. There are two likely explanations for this result. The first is that the duplicate sulfatases have essential roles that are of small, additive effect or are more visible in contexts other than a non-challenging environment. Alternatively, the genes may have become (or were upon arrival) functionally redundant, in which case they should be under relaxed selection in extant populations.

### Switch-gene paralogs show signatures of selection

Because the possible functions of the paralogs of *eud-1* remain unknown, we performed selection analyses to determine whether functions should be expected for these paralogs. To test the alternative hypothesis that the gene duplicates are undergoing nonfunctionalization, we examined patterns of selection on these genes in *P. pacificus.* Although mRNAs of all three sulfatase genes have been confirmed in the wild-type strain, expression itself does not equate to functionality of a product (Zheng et al. 2009; Kalyana-Sundaram et al. 2012). It is possible that gene duplicates, especially one born of the more recent duplication, *sul-2.2.1,* are dispensable and in the process of becoming pseudogenes. In this case, incipient pseudogenes should be under relaxed selection, showing similar rates of synonymous and nonsynonymous substitutions (Nei 1987). When we tested this possibility, we inferred the global *p*_N_/*p*_S_ = 0.125 (95% CI =[0.098, 0.156]) for *sul-2.1* and *p*_N_/*p*_S_=0.336 (95% CI=[0.244, 0.448]) for *sul-2.2.1* across populations of *P. pacificus.* The two duplicate sulfatase genes, therefore, showed signatures of purifying selection. These signatures were similar to that of *eud-1,* for which we inferred *p*_N_/*p*_S_=0.141 (95% CI=[0.095, 0.198]). Likewise, interspecific selection analyses showed the mean pairwise *d*_N_/*d*_S_=0.0665 (range=[0.0537, 0.0735]) for *sul-2.2.1* in *P. pacificus* and *P. exspectatus,* on the same order of magnitude as that estimated for *eud-1* (*d*_N_/*d*_S_=0.0523, range = [0.0464, 0.0588]). Thus, we rejected the hypothesis that the duplicate genes are neutrally evolving.

In addition to global *d*_N_/*d*_S_ signatures, we analyzed selection on individual codons to infer whether putative functional domains in the proteins have also been maintained by selection. In all three sulfatase genes, we identified several individual codons under purifying selection. In *sul-2.1,* 26 negatively selected codons were identified throughout the gene (Table S4). Fewer negatively selected codons were detected in *sul-2.2.1* (Table S5), although two of them encode residues (^86^Ser and ^90^Ala) in the conserved CSPSRA motif of the active site of the sulfatase. This motif is hypothesized to be particularly vulnerable in the protein, as coding mutations in its residues can eliminate enzymatic activity in homologs in humans (Litjens et al. 1996) as well as nematodes (Ragsdale et al. 2013). Likewise, codons for exactly the two homologous residues of EUD-1 (^96^Ser and ^100^Ala) were found to be under purifying selection (Table S6), suggesting *sul-2.2.1* specifically encodes a sulfatase maintained functional by selection. Additionally, the complete absence of null mutations in any extant haplotype supports the evolutionary maintenance of functional proteins across all populations of *P. pacificus.* Taken together, analyses of natural variation revealed signatures of strong purifying selection on both paralogs of the switch gene *eud-1.* Our findings support a historical scenario of functional specialization following serial duplications of an ancestral sulfatase gene: all three genes have necessary if unidentified functions, likely as sulfatases, and EUD-1 alone has assumed control of the mouth-polyphenism switch.

## Discussion

In this study we show that genomic evolutionary processes, particularly gene duplications, resulted in the specialization of a gene as a dosage-dependent operator of a polyphenism switch. We suggest that our findings affirm the utility of placing functional genetics approaches into a context of natural variation and the phylogenetic histories of genes and species. In such an approach, we have analyzed the history and phenotypic influence of genes that have undergone a lineage-specific radiation, particularly in organisms that have acquired a developmental novelty, morphological polyphenism. As a result we have shown that a history of gene duplications produced a polyphenism switch gene, thereby diverging from close paralogs that have other, unknown functions strongly maintained by selection.

### Duplication and specialization of a polyphenism switch-gene

The vast majority of duplicated genes are predicted to become nonfunctional (Haldane 1933; Nei and Roychoudhuri 1973; Lynch and Force, 2000). The process of gene silencing by relaxed selection, followed by fixation of deleterious alleles, is expected to happen rapidly, after which surviving duplicates may experience strong purifying selection (Lynch and Conery 2000). Here, we show that duplication events producing the switch gene *eud-1* did not result in neutral genomic expansion but rather in essential genes. Assays of general life-history traits were insufficient in this study to detect fitness disadvantages of mutants in any combination, other than the effect of *eud-1* on the mouth polyphenism, so the functions of those genes remain unknown. Nevertheless, it is possible that fitness consequences or even some influence on the mouth polyphenism are manifest in other, untested environmental contexts. Despite the limitations in determining phenotypes, selection analyses of *eud-1* paralogs indicate that the genes are experiencing purifying selection to maintain their functionality, presumably as sulfatases with a conserved active site.

We also found no qualitative evidence for a role of the other sulfatases, either singly or in combination, in dauer formation. Environmental stressors induce the Eu morph and dauer through inhibition of the same endocrine signaling module, the nuclear hormone receptor DAF-12 and its ligand Δ7–dafachronic acid (Bento et al. 2010). However, regulatory pathways for the mouth polyphenism and dauer formation do not completely overlap. For example, DAF-16/FOXO, which is an essential component of the dauer-formation cascade in both *C. elegans* and *P. pacificus*, has no role in the mouth polyphenism in the latter species (Ogawa et al. 2011). It is thus possible that ancestral sulfatases evolved roles independently of dauer induction or, alternatively, that one, two, or all of the sulfatases have only quantitative effects outside of the dauer-induction pathway as described for *C. elegans*(Hu 2005). The latter case is suggested by the link of sulfated sterols to dauer formation (Carroll et al. 2006), despite the unknown function or identity of the modified sterols. Given the lack of obvious mutant phenotypes, we predict that the other sulfatases play more nuanced roles in regulating phenotypic responses to the environment. Drawing on what is known of homologous human sulfatases, which play important roles in metabolism (Diez-Roux and Ballabio 2005), we are led to speculate a link of nematode sulfatases also to metabolic processes, whether dauer-relevant or otherwise. A more detailed understanding of the genetic context of EUD-1 will reveal whether hormonal or specific metabolic cues have been captured by a nutritionally responsive developmental decision.

Here, our phenotypic analyses of mutant paralogs in *P. pacificus* suggest that EUD-1 has specialized as a polyphenism regulator. If the paralogous sulfatases have other roles in the polyphenism, those roles must be distinct from that of the dosage-dependent switch activity of EUD-1. We hypothesize a scenario either of neofunctionalization, for example to accept or specialize on a new substrate, or of the division of roles from an ancestral enzyme (Jensen 1976; O'Brien and Herschlag 1999). Consequently, our study did not uncover evidence of degeneration and complementation for a single function. Evidence that selection has maintained the coding regions of *sul-2.1* and *sul-2.2.1* largely intact, including an essential motif in the active site of *sul-2.2.1,* suggests that at least the catalytic domain is functional in all daughter sulfatases. Given that these genes likely encode functional sulfatases, the enzymes may have diverged to assume different substrate specificities, such as by changes in domains outside of the catalytic core. Alternatively, the genes may have diverged in expression in time or space, either in transcriptional regulation or in posttranslational transport. The former possibility is an obvious one, and although potential promoter sequences have already diverged beyond reliable alignment, experimental mapping of regulatory elements could detect causal differences of gene expression. In the case of posttranslational differences, diverged expression could result from the less conserved coding sequences of *eud-1* and *sul-2.2.1.* For example, the first exon of *eud-1* encodes part of a predicted signal peptide possibly marking the protein for cellular export (Fig. 2), but the sequence of this exon is missing at the locus of *sul-2.2.1* in *P. pacificus* and *P. exspectatus.* The question of whether idiosyncratic coding motifs such as this are functionally important, or simply neutral gains or losses after duplication, remains for experimentation.

Despite the evidence for fixation of duplicate genes, at least one of specialized function, how the duplications survived mutational degeneration before diverging still must be explained (Bergthorsson et al. 2007). A speculative possibility is the immediate fitness advantages of gene amplification, allowing selection for higher enzyme dosage (Mouches et al. 1986; Kondrashov and Kondrashov 2006). Gene dosage is known to be a defining feature of EUD-1 activity, where increased dosage results in a higher frequency of the Eu morph (Ragsdale et al. 2013). If functions of an ancestral sulfatase included a role in the switch, conditions favoring greater sensitivity of Eu development could select for amplification of the enzyme prior to functional specialization. Given a sufficiently large population size (i.e., 10^6^), which is possible in amphimictic nematodes (Dey et al. 2013), the fixation of non-complementary duplicates with a selective advantage would be possible (Lynch et al. 2001; Walsh 2003). Extending functional studies to other species could test the possibility of amplification, by determining whether other species with duplicated genes show additive effects on the polyphenism. The new feasibility of reverse genetics offers the potential to scale up comparative functional tests of plasticity evolution.

### Reverse genetics to study evolution by gene duplication

Beyond offering insights into history of a polyphenism, our study gives a tractable example of a universal genetic technique to study evolution by gene duplication. The use of CRISPR/Cas9 for cost-effective gene knockouts in untraditional model systems promises to revolutionize the mechanistic study of ecology and evolution (Bono et al. 2015). Obvious targets have included genes suggested by transcriptomic studies or functions in other organisms, and here we explore genes closely related to factors revealed in our study organism by forward genetics. With the ability to study multiple candidate genes in the same individual, the effects of gene duplication on phenotypes can add a new dimension to functional genetics (Wang et al. 2015), and it is known that sgRNAs can delete multiple targets simultaneously (Endo et al. 2015; Peng et al. 2015; Prykozhij et al. 2015). In combination with simple genetic crosses, we have likewise used multiple gene deletions to distinguish among historical scenarios following gene duplications. Despite the obvious pitfall of unknown off-target effects (Hsu et al. 2013), particularly on genes with identical sequence regions, the concurrent advent of genomics in non-model organisms (Ellegren 2014) increasingly reduces the liability of unknown sequences. Therefore, we suggest our study design might be more broadly applied to questions of evolution by gene duplication in other systems.

## Acknowledgments

This work was funded in part by the National Science Foundation (NSF IOS-1557873).

